# Early root architectural traits and their relationship with yield in *Ipomoea batatas* L

**DOI:** 10.1101/2023.10.20.563294

**Authors:** L.O. Duque, G. Hoffman, K. Pecota, G. C. Yencho

**Affiliations:** . Department of Plant Science, The Pennsylvania State University, University Park, PA 16802, U.S.A.; . Department of Food Science, The Pennsylvania State University, University Park, PA 16802, U.S.A.; . Department of Horticultural Science, North Carolina State University, Raleigh, NC 27695, U.S.A.

## Abstract

Root system architecture in storage root crops are an important component of plant growth and yield performance that has received little attention by researchers because of the inherent difficulties posed by *in-situ* root observation. Sweetpotato (*Ipomoea batatas* L.) is an important climate-resilient storage root crop of worldwide importance for both tropical and temperate regions, and identifying genotypes with advantageous root phenotypes and improved root architecture to facilitate breeding for improved storage root yield and quality characteristics in both high and low input scenarios would be beneficial. We evaluated 38 diverse sweetpotato genotypes for early root architectural traits and correlated a subset of these with storage root yield. Early root architectural traits were scanned and digitized using the RhizoVision Explorer software system. Significant genotypic variation was detected for all early root traits including root mass, total root length, root volume, root area and root length by diameter classes. Based on the values of total root length, we separated the 38 genotypes into three root sizes (small, medium, and large). Principal component analysis identified four clusters, primarily defined by shoot mass, root volume, root area, root mass, total root length and root length by diameter class. Average total and marketable yield and number of storage roots, was assessed on a subset of eight genotypes in the field. Several early root traits were positively correlated with total yield, marketable yield, and number of storage roots. These results suggest that root traits, particularly total root length and root mass could improve yield potential and should be incorporated into sweetpotato ideotypes. To help increase sweetpotato performance in challenging environments, breeding efforts may benefit through the incorporation of early root phenotyping using the idea of integrated root phenotypes.

## 1. Introduction

Plant roots perform several fundamental roles such as, anchorage, nutrient and water procurement followed by the transport of these to the stem, storage of reserves in the form of non-structural carbohydrates, and environmental sensing of edaphic abiotic and biotic constituents (Aubrey and Teskey, 2018; Ortiz-Castro and Lopez-Bucio, 2019; Stubbs et al., 2019; Lynch et al., 2021), which ultimately impact plant fitness, adaptation, and productivity (Shekhar et al., 2019; Canto et al., 2020; Griffiths and York, 2020). Root system architecture is important in natural and managed ecosystems (i.e., forests, pastures, and rangelands), but particularly pertinent to crop production systems, as they are more intensively managed with added inputs (i.e., agrochemicals and irrigation) and improved seed making the latter indispensable for human well-being and progress (Lynch, 2022). Crop growth and yield depend on the ability of the root system to explore, forage, and secure edaphic resources (Qiao et al., 2018; Chen et al., 2022; Jorda et al., 2022). However, root phenotyping research has lagged behind its aboveground counterpart due to the challenging nature of direct selection of root traits under field conditions, which because root systems are underground, requires significant logistical and infrastructure costs, time, and management considerations (Smith and De Smet, 2012; Thomas et al., 2016). Additionally, selecting directly for root traits is often difficult due to a lack of phenotypic diversity resulting from domestication and/or a critically reduced population size, limiting our ability to broadly survey the genetic diversity of a crop species (Bus et al., 2011; Thomas et al., 2016) or in other cases, diverse genetic material is not straightforwardly tested in the field due to being unadapted to local environmental conditions (Thomas et al., 2016).

Direct phenotypic selection for storage root traits and yield is the *rule-of-practice* in many storage root and tuber crops such as cassava [*Manihot esculenta* Crantz; (Sunvittayakul et al., 2022)], sweetpotato [*Ipomoea batatas* L., (Shumbusha et al., 2017)], and potato [*Solanum tuberosum* L., (Dolnicar, 2021)]. However, when storage roots and tubers of these crops are harvested, many if not all of the *adventitious roots* (AR) [i.e., roots that are formed from any non-storage or root tissue and are produced during normal development (e.g., crown roots on maize and nodal roots on sweetpotato) or in response to an abiotic and/or biotic stress condition] are lost and not phenotyped. To circumvent these issues, many high throughput phenotyping (HTP) root screening systems have been developed, used, and scrutinized by the research community (Thomas et al., 2016; Tracy et al., 2019). However, results from these HTP root system studies are highly variable and require a large number of replicates to assess sources of variation (Adu et al., 2014). In principle, such root HTP systems are not capable of the thorough evaluation of soil borne roots. They can, however, identify early differing root traits that can be then compared to their mature counterparts in the field (Thomas et al., 2016). To date, there are just a handful of phenotyping studies that have been designed to examine the associations between early root growth and field performance (White et al., 2013; Ruiz et al., 2018; Strock et al., 2019). Early AR phenotypes may have significant effects on fitness and adaptation and are more easily measured than their mature counterparts. Early root phenotyping is now a common practice and it has been demonstrated to be an effective method for characterizing the root genetic diversity and phenotypic plasticity among important crop species including rice (Tomita et al., 2017), maize (Sanchez et al., 2018), wheat (Ruiz et al., 2018), pearl millet (Passot et al., 2016), common bean (Lynch and van Beem, 1993), and sorghum (Joshi et al., 2017). In root and tuber crops (RTC), early root phenotyping is now an emerging field and has revealed important advances in sweetpotato (Villordon et al., 2020), cassava (Selvaraj et al., 2019), and potato (Tiwari et al., 2020).

Sweetpotato, is one of the most important staple food crops in developing countries worldwide, where it ranks six overall after maize, rice, wheat, potatoes, and cassava in terms annual production (FAO, 2022). Sweetpotato produces a starchy storage root that is used to prepare a wide variety of foods and other products, and like many other vegetable commodities, are marketed in several ways depending on industry and consumer demand. Sweetpotatoes are usually produced for fresh consumption but now are progressively used for alternate markets, such as processed foods (e.g., French fries and chips) and commercial products (e.g., starch, flour, and food dyes) and suchlike (Duque et al., 2022). Additionally, sweetpotato’s wide climatic adaptation and use in a variety of cropping systems is thought to be related to its perennial nature and it’s abiotic tolerance, all of which have fundamental implications for meeting nutritional requirements, reducing poverty, and alleviating food insecurity (El Sheikha and Ray, 2017). Vine cuttings from sprouted sweetpotatoes (i.e., “slips”) are used as propagation and/or planting material. The storage roots developed from these are AR and normally arise from the underground stem portion (Togari, 1950; Belehu et al., 2004). According to Togari (1950) and (Kays, 1985) the adventitious roots of sweetpotato are subdivided into ‘thick’ and ‘thin’ roots, which are further differentiated into fibrous, ‘pencil’, and storage roots. Environmental (e.g., air and soil temperature), edaphic characteristics (e.g., soil physical properties and fertility), and genotype influence the formation and growth of storage roots (Ravi and Indira, 1999). Hence, an adapted genotype and favorable growing conditions during the transition from primary to secondary root development have a direct effect on the onset of storage root formation, which is marked by the activity of the vascular cambium, and anomalous primary and secondary cambial elements (Wilson, 1971; Wilson, 1982; Kays, 1985).

In sweetpotato, root traits have only recently garnered attention (Villordon et al., 2014; Duque and Villordon, 2019). Only a few studies have characterized sweetpotato AR development under controlled environmental settings, for example, lateral root length, count, and surface area under water stress (Pardales and Yamauchi, 2003; Villordon et al., 2012), lateral root development under reduced nitrogen and phosphorus availability (Villordon et al., 2013; Villordon et al., 2018). Measuring the impact of individual root architectural traits under edaphic stress warrants a more thorough understanding of root system function that can be then implemented in an ‘ideotype breeding’ strategy (York et al., 2013; Lynch, 2015). Thus, linking early root system architecture and traits to storage root formation under the direct or indirect influence of biotic and abiotic interactions in sweetpotato is of paramount importance for understanding how these are conducive to storage root yield potential.

However, care is to be taken when analyzing a set of elite sweetpotato germplasm with potentially limited root phenotypic variation. This situation can provide an inaccurate assessment of the actual genetic variation available for any given trait, but also masks the ability to identify the effects of root system variation on plant performance. In sweetpotato, there exists limited knowledge in root phenotyping of diverse germplasm, hence, examining the root system architecture as a whole and connecting these early root traits to yield performance in the field are conducive to understanding the linkage between sweetpotato germplasm diversity and its growing environment. To our knowledge, there are no published studies on early adventitious root traits in sweetpotato and its relationship to yield across different environments.

This study used a semi-hydroponic ebb and flow phenotyping system to characterize root traits of 38 sweetpotato genotypes grown in different parts of the world. We hypothesize that significant variation exists in early root traits within the germplasm tested and that cultivars differ in traits in contrasting environments. This potential variation in early root traits can partially explain both water and nutrient acquisition and influences overall root function. We conducted studies to determine if early root traits measured on adventitious roots of clonally propagated sweetpotato were correlated to key storage root characteristics in the field. The results of these studies have broadened our understanding of the comparative involvement of early individual root traits of a neglected storage root crop, and may aid in the selection of sweetpotato genotypes with differing root architectures and performances for further studies in both controlled greenhouse and field environments.

## 2. Materials and Methods

### 2.1 Sweetpotato genotypes

A collection of 38 sweetpotato [(*Ipomoea batatas* (L.) Lam.] genotypes including cultivars, breeding lines, and cultivated clones originating from seven broad regional groups: Africa, the Caribbean, Central America, the Far East, North America, the Pacific Islands, and South America, were studied (Table 1). All genotypes were obtained courtesy of the North Carolina State University Sweetpotato and Potato Breeding and Genetics Program, Raleigh, NC. The diversity panel subset from North America reflects the trend of commercial sweetpotato cultivars commonly sowed throughout the United States, while the rest of broad regional groups represents sweetpotato germplasm adapted to those specific centers of origin (Table 1).

**Table 1.**
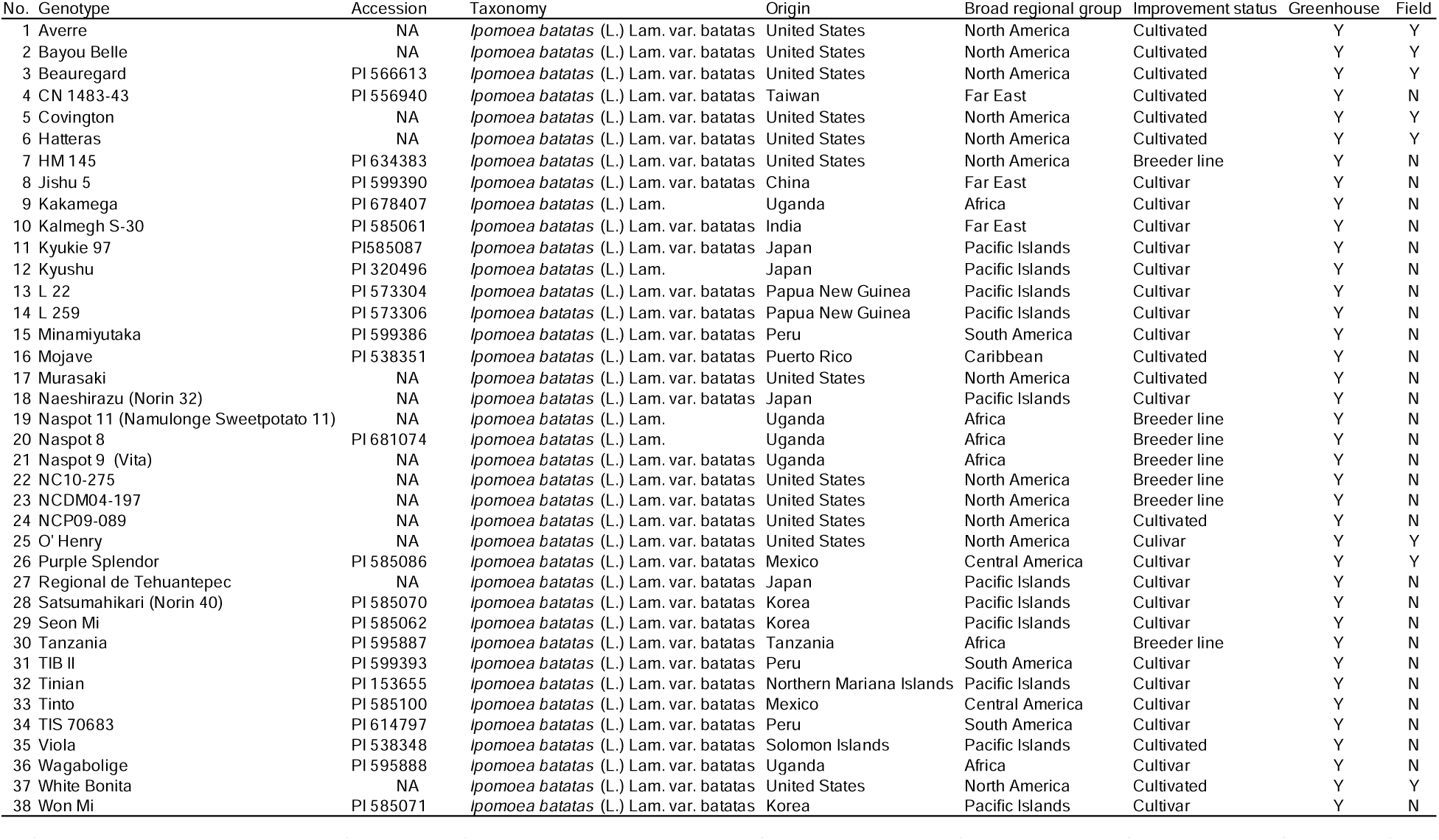
List of 38 sweetpotato genotypes evaluated for root traits using a semi-hydroponic ebb and flow phenotyping system in the greenhouse at Penn State University and field experiments at Rock Springs, PA, USA in 2022. A subset of eight genotypes were used for field phenotyping of yield traits. All sweetpotato genotypes were provided by North Carolina State University.

### 2.2 Greenhouse experimental design and growth conditions

The phenotyping experiment was conducted in a greenhouse under controlled conditions at the Pennsylvania State University, University Park, PA, USA (40° 85′ N, 77° 82′ W), from April to May 2022 using an expandable ebb and flow system semi-hydroponic phenotyping system (Figure 1). Four sweetpotato stem cuttings (‘slips’) of each genotype were planted in five replicate phenotyping platforms arranged in a randomized complete block design with one phenotyping platform as one replicate. Each phenotyping platform could accommodate up to 12 stem cuttings distributed in 12 independent growth units. Buffer stems were used when necessary to guarantee equal number of stems per phenotyping platform.

**Figure 1.**
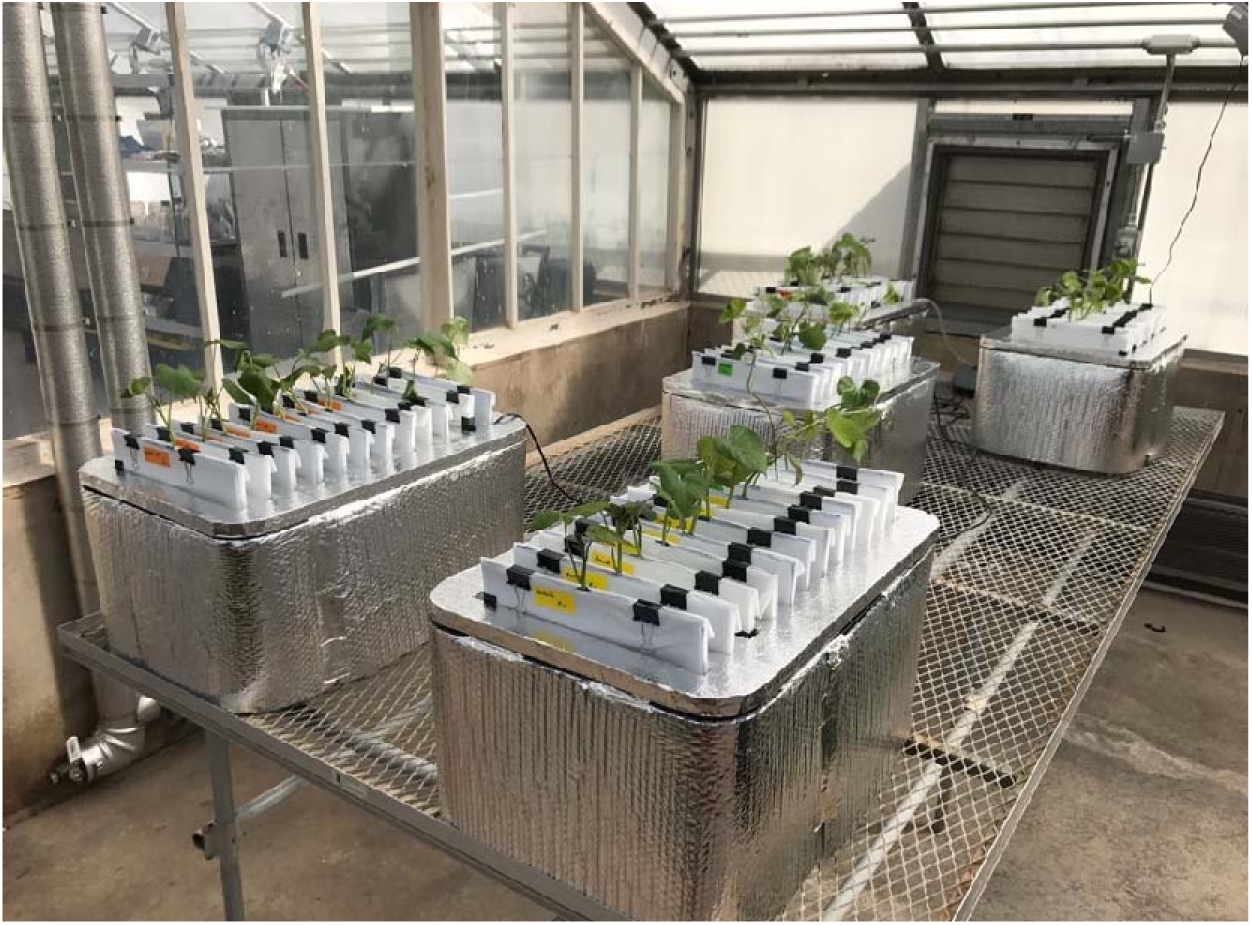
Sweetpotato semi-hydroponic ebb-and-flow soilless phenotyping platform constructed and tested for stem-derived adventitious roots. The root phenotyping experiment was conducted in a greenhouse located at Penn State University in 2022.

The details and description of the semi-hydroponic system are described in Duque (2023) with modifications as follows. The watering system was filled with 180 L of full-strength Peters 5-11-26 hydroponic nutrient solution (Peters 5-11-26 Hydroponic Special Fertilizer, Everris NA Inc., Dublin, OH, USA) supplemented with Mg (2.46 mM; Epsoak Agricultural Epsom Salt, SF Slat Co., Alameda, CA, USA), N, and Ca (7.14 mM N, 2.49 mM Ca; calcium nitrate, Jack’s Nutrients 15-0-0 Cal Nit Part B, JR Peters Inc., Allentown, PA, USA). The concentrations of the macro/micro-nutrients present in the hydroponic nutrient solution are provided in Table 2. The nutrient solution stored in the nutrient supply tank was delivered to the phenotyping platform via an automatic submersible pump through a time controller. The periodic pumping was set as 10 min on and 240 min off during a 24-hour period. The nutrient solution was refreshed weekly. Greenhouse environmental growth conditions exhibited a photoperiod of 14/10 h at 32/28 °C (light/darkness) with a maximum midday photosynthetic flux density of 1200 µmol photons m−2 supplemented with LED lights. The ambient humidity was 40%.

**Table 2.** Composition of macro and micronutrients in the liquid media used in the greenhouse study.

| Product | Element | Concentration (ppm) |
| --- | --- | --- |
| Peters Professional 5-11-26 | Nitrogen (all nitrate N) | 50 |
| Hydroponic Special Fertilizer Part A | Phosphorus (P) | 48 |
|  | Potassium (K) | 216 |
|  | Magnesium (Mg) | 60 |
|  | Sulfur (S) | 80 |
|  | Iron (Fe) | 3 |
|  | Manganese (Mn) | 0.5 |
|  | Zinc (Zn) | 0.15 |
|  | Copper (Cu) | 0.15 |
|  | Boron (B) | 0.5 |
|  | Molybdenum (Mo) | 0.1 |
| Epsoak Agricultural Epsom Salt | Magnesium (Mg) | 296.1 |
| Jack's Nutrients Boost 15-0-0 | Nitrogen (as nitrate; $\text{NO}_3$) | 93 |
| Cal Nit Part B | Nitrogen (as ammonium; $\text{NH}_4$) | 7 |
|  | Calcium (Ca) | 116.1 |

### 2.3 Field experiment

The field experiment was conducted in 2022 at the Russell E. Larson Agricultural Research Center of The Pennsylvania State University located in Rock Springs, PA (lat. 40°42’38.1”N, long. 77°57’52.07”W). The field site is dominated by a Murrill channery silt loam soil (fine-loamy, mixed, semiactive, mesic Typic Hapludult), and slopes from 0% to 3%. Monthly temperature and precipitation data were obtained from the Network for Environmental and Weather Applications, Rock Springs, PA, weather station (lat. 40°42’32.5”N, long. 77°57’06.9”W). All sweetpotato slips used for both years were G2 planting stock provided by North Carolina State University, Raleigh (Table 1). A combination raised bed–mulch–drip tape layer (model 2400 Mini Layer; Rain-Flo Irrigation, East Earl, PA) was used to form raised beds 91 cm wide and 20 inches high. Each bed was covered with 1-mil thick black embossed plastic mulch (part no. BLK324, Rain-Flo Irrigation). Distance between bed centers was 91 cm. A single line of 6-mil-thick drip irrigation tape with 20-cm emitter spacing (T-Tape; T Systems International, San Diego, CA) was laid under the plastic mulch for each row. Slips were spaced 30 cm apart in a single row and hand-planted into plastic mulch on 5 June 2022. All trials used a randomized complete block design with four replicates of 10 plants per experimental plot, with a 91 cm break between plots. Soil tests were done 15 d before field preparation and were performed by the Agricultural Analytical Services Laboratory of The Pennsylvania State University, University Park. Soil tests showed phosphorus (P) levels at 239 kg Ha^-1^ and potassium (K) levels at 221 kg Ha^-1^ as determined by the Mehlich 3 soil test. Before laying down the black plastic mulch, 39 kg Ha^-1^ of calcium nitrate (15.5N–0P–0K) and 61 kg Ha^-1^ of magnesium sulfate (0N–0P– 0K–9Mg) were incorporated preplanting based on soil test reports. After 30 d (storage root initiation period), an additional 39 kg Ha^-1^calcium nitrate was fertigated. The fertilization regime remained the same for both years. All trials were drip irrigated as needed using the ‘‘feel and appearance’’ method (USDA, 1998) throughout the growing season. Manual weeding between rows was done periodically in June and July. By August, sweetpotato vines grew and spread enough to prevent any weed growth between the rows. Sweetpotatoes were harvested on 5 Oct. 2022, which corresponded to 120 d after transplanting. Vines were detached using a walk-behind string trimmer (model ST 100; Cub Cadet, Cleveland, OH), and plastic mulch was removed by hand. Storage roots were lifted using a one-row potato digger (model D-10M; U.S. Small Farm Equipment Co., Worland, WY) and were sorted into the following categories using U.S. standards for grades of sweetpotatoes (USDA-AMS, 2005). After the storage roots were categorized, they were weighed to calculate marketable yield (MY) and total yield (TY), where MY equals the sum of all categories excluding cull and TY equals MY plus cull. Total yield (TY), marketable yield (MY), and number of storage roots (SR#) were calculated for the field experiment.

### 2.4 Greenhouse root data collection and image analysis

Roots from all genotypes were harvested after 15 days and root section samples were excised and scanned in grayscale at 600 dpi using a desktop scanner (Epson Perfection V800, Long Beach, CA, USA), and root images for each 5 cm sections proximal to the stem were analyzed using RhizoVision Explorer v2.0.3 (Seethepalli et al., 2021), where the image thresholding level was set at 200 pixel intensity, noisy component filtering set at 0.2 mm^2^, and root pruning threshold was set at 1 pixel. A root diameter threshold of 0.2 mm was used to distinguished adventitious roots from 1^st^ order roots. Root morphology data, such as, number of root tips, total root length, root area perimeter, average root diameter, root volume, surface root area, and root branching frequency, were generated in the RhizoVision Explorer program (Table 3). The following traits were then calculated from the measured data following Qiao et al. (2019):

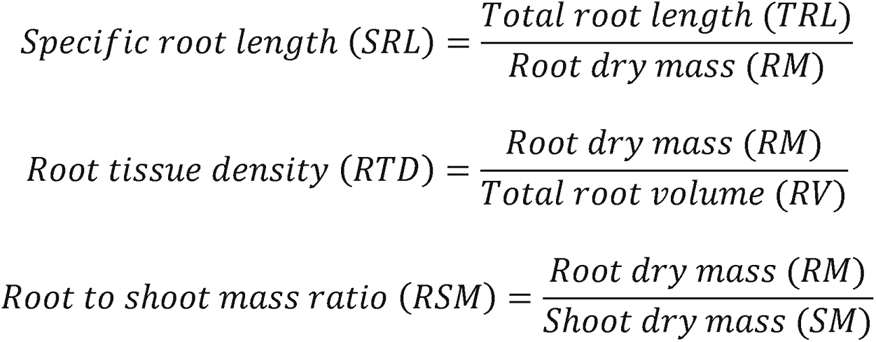

**Table 3.** Description of traits measured for both greenhouse and field experiments. Thirty-eight sweetpotato genotypes were grown in a semi-hydroponic ebb-and-flow soilless phenotyping platform at Penn State University. Eight sweetpotato genotypes were grown in the field at Rock Springs, PA.

| Traits | Abbreviation | Description | Units | Experiment |
| --- | --- | --- | --- | --- |
| Roots in node 1 | RN1 | Total number of roots in node 1 per plant | count | greenhouse |
| Roots in node 2 | RN2 | Total number of roots in node 2 per plant | count | greenhouse |
| Roots in node 3 | RN3 | Total number of roots in node 3 per plant | count | greenhouse |
| Shoot mass | SM | Total shoot dry mass per plant | mg | greenhouse |
| Root mass | RM | Total root dry mass per plant | mg | greenhouse |
| Specific root length | SRL | Total root length divided by root dry mass | cm mg<sup>-1</sup> | greenhouse |
| Root tissue density | RTD | Total root dry mass divided by root volume | mg mm<sup>3</sup> | greenhouse |
| Root shoot mass ratio | RSM | Total root dry mass divided by shoot dry mass | - | greenhouse |
| Root tips | RT | Total number of root tips per plant | count | greenhouse |
| Root length | TRL | Total length of all roots per plant | cm | greenhouse |
| Branching density | BD | Total number of branch points divided by the total root length | branch cm<sup>-1</sup> | greenhouse |
| Average root diameter | RD | Average diameter of all roots per plant | mm | greenhouse |
| Root volume | RV | Total volume of all roots per plant | cm<sup>3</sup> | greenhouse |
| Root area | RA | Total surface area of all roots per plant | cm<sup>2</sup> | greenhouse |
| Root length in diameter class 1 | RLD1 | Root length of roots in diameter class < 0.5 mm | cm | greenhouse |
| Root length in diameter class 2 | RLD2 | Root length of roots in diameter class 0.5 to 1.0 mm | cm | greenhouse |
| Root length in diameter class 3 | RLD3 | Root length of roots in diameter class ≥ 1.0 mm | cm | greenhouse |
| Total yield | TY | Total fresh weight of U.S. No.1, jumbos, canners and culls storage roots categories | kg Ha<sup>-1</sup> | field |
| Marketable yield | MY | Total fresh weight of U.S. No.1, jumbos, and canners storage roots categories | kg Ha<sup>-1</sup> | field |
| Number of storage roots | SR# | Total number of mature storage roots in field | count | field |

A total of 16 root traits and one shoot trait were measured for the greenhouse experiment and three traits were measured for the field experiment (Table 3).

### 2.5 Statistical analysis

Data were tested for normality and subjected to analysis of variance using JMP Pro (version 16; SAS Institute, Cary, NC) on all measured root traits. When comparing genotypes and root traits, if the overall F test was significant (*p* ≤ 0.05), means separations were evaluated using Tukey’s Honest Significant Difference test at *p* ≤ 0.05. Pearson’s correlations coefficients were calculated for all greenhouse and field root traits and were considered statistically significant at *p*_≤_0.05. Traits with coefficient of variation (CV) values ≥_0.3 were selected for principal component analysis to identify the determinants of root architecture variability across genotypes. Principal component analysis (PCA) was implemented to discover root traits and yield characteristics contributing to the observed variation among the genotypes, and to identify correlated variables. To avoid potential differences in the units of measurements, all data were centered and scaled before the PCA analysis.

## 3. Results

### 3.1 Summary statistics for root traits of sweetpotato genotypes

Root traits revealed significant differences in the pattern of root development and distribution, and the majority differed among sweetpotato genotypes (Table 4). There were wide ranges in values and variations for all root traits measured such as RM (4.7 – 1092.8 mg; CV = 0.96), RT (29.3 – 39658.3; CV = 1.68), TRL (65.5 – 10815 cm; CV = 0.83), RBD (0.8 – 51.2 branch cm^-1^; CV = 0.76), RA (78.5 – 24079 mm^2^; CV = 0.98), and RV (7.1 – 10056.9 mm^3^; CV = 1.51), were relatively high (all *p* ≤ 0.0001), among others. SM measured at harvest (133.1 – 6480.3 mg; CV = 0.63), differed significantly among genotypes tested (all *p* ≤ 0.0001) (Table 4).

**Table 4.** Descriptive statistics of 17 measured traits in 38 sweetpotato genotypes grown in a semi-hydroponic ebb-and-flow soilless phenotyping platform.

| Traits | Min | Max | Mean | Median | Std Dev | CV | p value |
| --- | --- | --- | --- | --- | --- | --- | --- |
| RN1 | 2 | 25 | 7.7 | 7 | 4.1 | 0.53 | 0.0001 |
| RN2 | 3 | 41 | 10.2 | 9 | 5.4 | 0.52 | 0.0001 |
| RN3 | 2 | 33 | 13.6 | 13 | 6.3 | 0.46 | <.0001 |
| SM | 133.1 | 6480.3 | 1470.3 | 1257.1 | 930.0 | 0.63 | <.0001 |
| RM | 4.7 | 1092.8 | 178.5 | 136.0 | 171.3 | 0.96 | <.0001 |
| SRL | 13.1 | 470.2 | 143.2 | 137.9 | 48.8 | 0.34 | <.0001 |
| RTD | 0.010 | 0.266 | 0.035 | 0.031 | 0.022 | 0.64 | 0.1601 |
| RSM | 0.01 | 0.41 | 0.11 | 0.10 | 0.06 | 0.58 | <.0001 |
| RT | 29.3 | 39658.3 | 2516.7 | 1546.6 | 4216.6 | 1.68 | <.0001 |
| TRL | 65.5 | 10815 | 2310 | 1855.3 | 1918.3 | 0.83 | 0.0001 |
| RBD | 0.8 | 51.2 | 9 | 7.3 | 6.8 | 0.76 | <.0001 |
| RD | 0.3 | 3.9 | 1.6 | 1.5 | 0.7 | 0.47 | 0.0362 |
| RV | 7.1 | 10056.9 | 729.7 | 405.6 | 1101.5 | 1.51 | <.0001 |
| RA | 78.5 | 24079.0 | 3230.9 | 2352.3 | 3175.1 | 0.98 | <.0001 |
| RLD1 | 18.4 | 79817.6 | 16978.3 | 12905.3 | 14214.7 | 0.84 | 0.0001 |
| RLD2 | 108.4 | 17814.0 | 3870.9 | 3145.8 | 2883.6 | 0.74 | 0.0235 |
| RLD3 | 0.0 | 23559.9 | 2251.1 | 1222.9 | 3059.4 | 1.36 | <.0001 |

RM ranged from 4.7 (Beauregard) to 1092.8 (NCDM04-197) mg per plant with a median value of 136 mg (Table 4). About 31.6% of the 38 genotypes had root mass values > 178.5 mg per plant. TRL ranged from 65.5 (Beauregard) to 10815 (Seon Mi) cm per plant with a median value of 1855.3 cm (Table 2). Among the 38 genotypes, seven genotypes had TRL < 1000 cm per plant, 24 genotypes had TRL values from 3000 to 1000 cm per plant, and seven genotypes had RL > 3000 cm (Figure 2). Based on the TRL value of each genotype compared to the mean value (2310 cm per plant) of TRL ± three standard error, the 38 sweetpotato genotypes were divided into three root sizes; 14 genotypes were considered as having a small root system (TRL < 17000 mm per plant), 14 genotypes were considered as having a large root system (TRL > 29000 mm per plant), and 10 genotypes were considered as having a medium root system (17000 ≤ TRL ≤ 29000 mm per plant) (Figure 2). The root mass, root length distribution of roots by diameter class (fine roots = diameter class < 0.5 mm; medium roots = diameter class 0.5 ≤ X ≤ 1.0; thick roots = diameter class ≥ 1.0 mm), followed similar trends to RL for the three root size classes (Figure 3). For example, root length by diameter class data showed that sweetpotato plants had relatively more fine roots with a mean root diameter of < 0.5 mm compared to medium and large roots for all root size categories (Figure 3C). Root length (fine roots) accounted for 73.5% of the total root length, while medium and large roots accounted for 16.7 and 9.7%, respectively.

**Figure 2.**
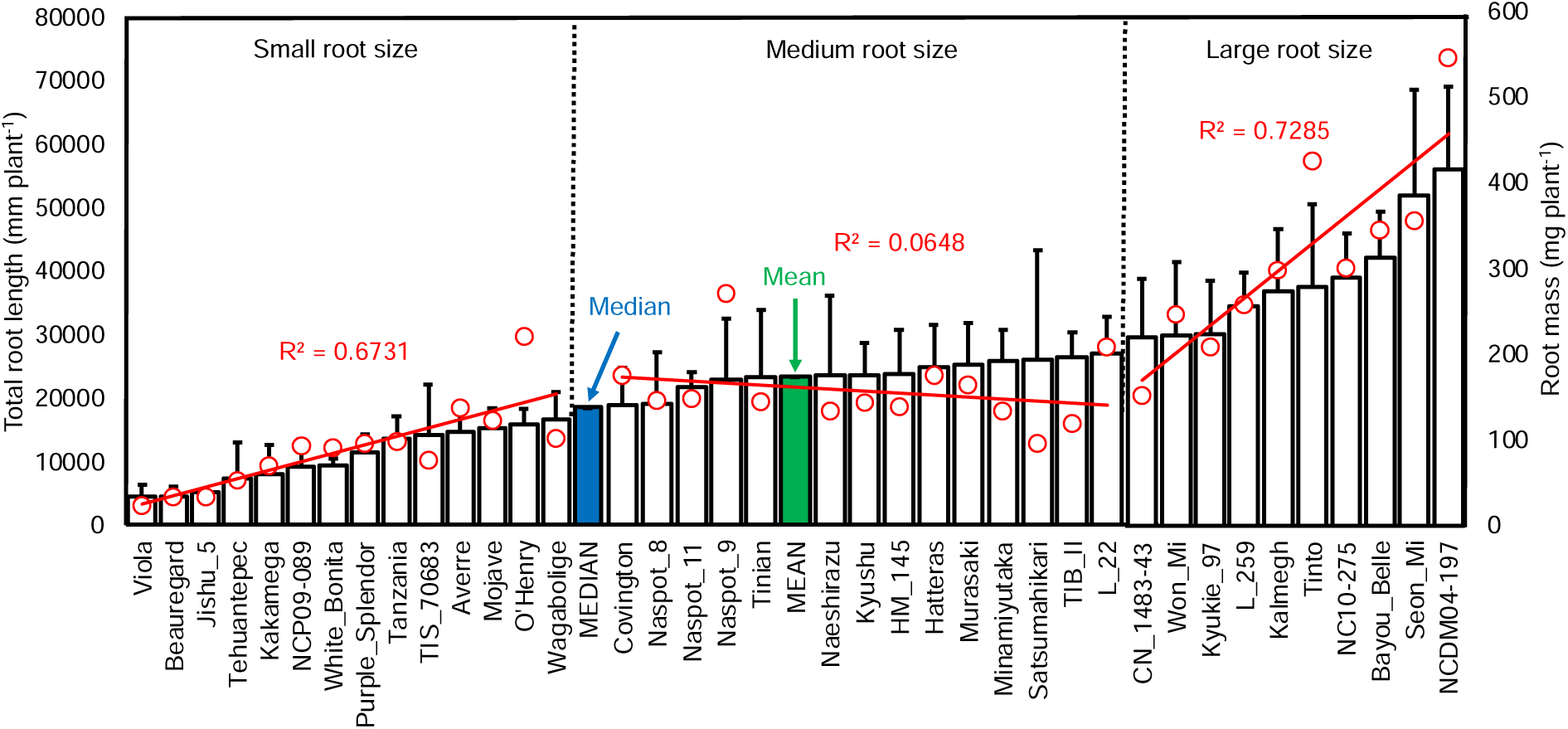
Vertical bar plot (ordered from shortest to longest mean total root length) showing genotypic variation for total root length and root mass of 38 sweetpotato grown in a semi-hydroponic ebb-and-flow soilless phenotyping platform 15 days after sowing. Blue bar indicates median value and green bar indicates mean value for total root length and plotted for all genotypes. Means, trendlines with correlation values (*R^2^*) for root mass for each genotype root size are given in red.

**Figure 3.**
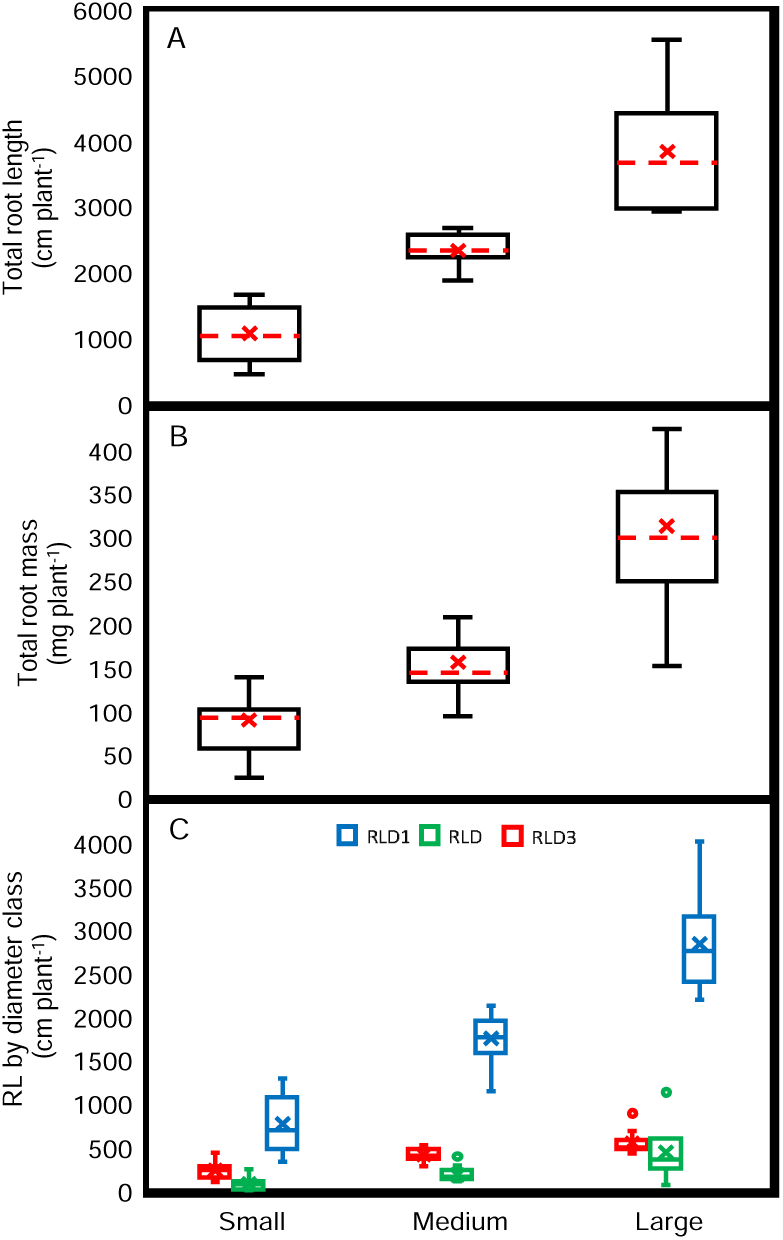
Total root length (A), total root mass (B), and root length by diameter class (C) for sweetpotato genotypes in three root size categories, small (14 genotypes), medium (14 genotypes), and large (10 genotypes). The 38 genotypes were grown in a semi-hydroponic ebb-and-flow soilless phenotyping platform 15 days after sowing. Significance differences are shown for the three root size categories (*p* < 0.05). For (A) and (B), box plot components show whiskers and box and are determined by the 5^th^ and 95^th^ percentiles, and 25^th^ and 75^th^ percentiles, respectively. The red dashed line and X inside the box mark the median and mean, respectively. For (C), blue boxplot (RLD1) indicates fine roots (diameter class < 0.5 mm), green boxplot (RLD2) indicates medium roots (diameter class 0.5 ≤ X ≤ 1.0 mm), red boxplot (RLD3) indicates thick roots (diameter class ≥ 1.0 mm), respectively.

### 3.2 Correlations among root traits

Pearson’s correlation coefficient analysis displayed strong correlations among several root traits (e.g., root mass, root tips, total root length, root diameter, root volume, root area, root volume, and root length in diameter class) (Table 5). In particular, the TRL was highly correlated with RM (r_=_0.83), SM (r = 0.80), RV (r = 0.75), RA (r = 0.93), and all three RLD measured branch number (0.90), root volume (0.85) and branch density (0.84) (each at *p* < 0.0001). BD was correlated with RD (r = 0.53), and RT (r = 0.77) (each at *p* < 0.0001). Interestingly, SRL showed significant negative correlations when compared to RM (r = -0.30; *p* < 0.0001), and RV (r = -0.18; *p* < 0.05). RTD was correlated with decreasing TRL (r = -0.32), RA (r = -0.34), RLD1 (r = -0.3), RLD2 (r = - 0.31), and RLD3 (r = -0.32) (each at *p* < 0.0001), and RV (r = -0.3; *p* < 0.05) (Table 5). Specifically, SM was positively correlated with both TRL (R^2^ = 0.65) and RM (R^2^ = 0.71) (Figure 4A). Also, TRL was positively correlated with RLD1 (R^2^ = 0.97), RLD2 (R^2^ = 0.83), and RLD3 (R^2^ = 0.68) (Figure 4B).

**Figure 4.**
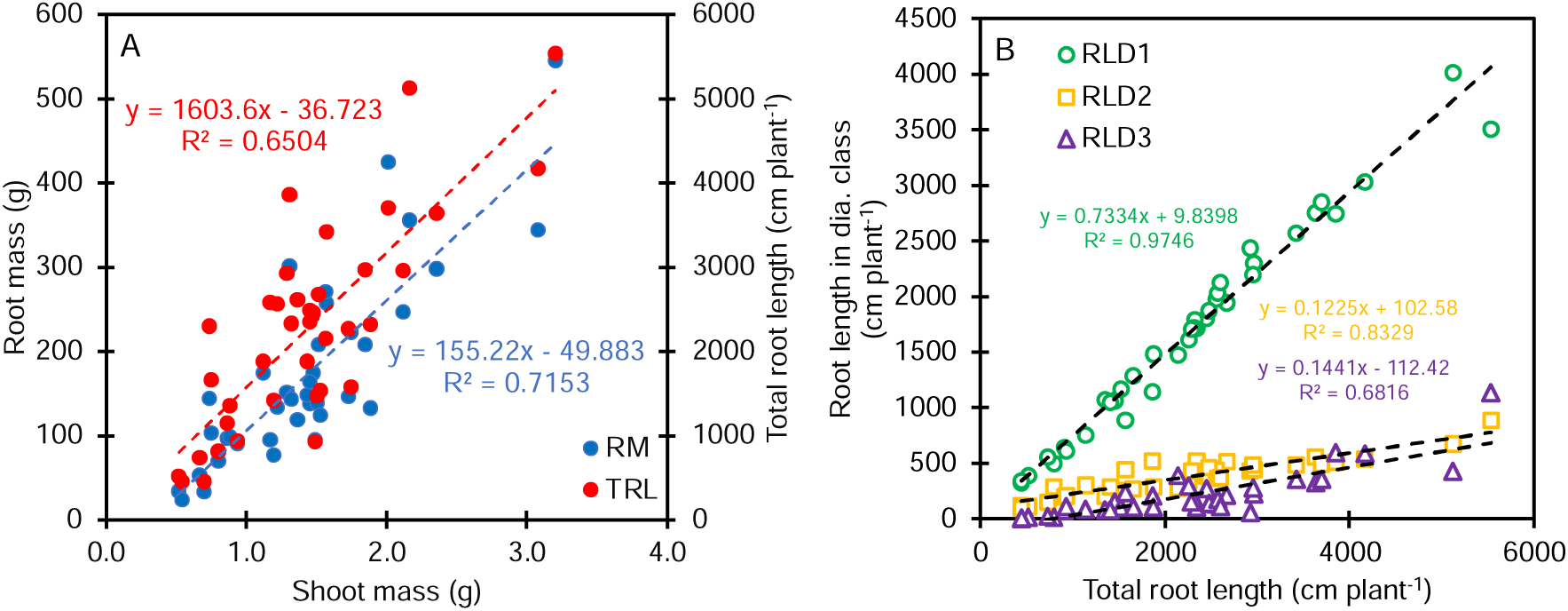
Correlations between (A) shoot mass, root mass, and total root length, (B) total root length, and root length in diameter class 1, 2, and 3. In 38 sweetpotato genotypes grown in a semi-hydroponic ebb-and-flow soilless phenotyping platform 15 days after sowing. For (A), red color dots correlates SM and TRL, blue color dots correlates SM and RM. For (B), green color open circles correlate TRL and RLD1, yellow open squares correlate TRL and RLD2, and purple open triangles correlate TRL and RLD3. Each dot and open symbol represent the mean values of four replicates per sweetpotato genotype.

**Table 5.** Pearson’s correlation matrix for 16 root traits and one shoot trait (SM) in 38 sweetpotato genotypes grown in a semi-hydroponic ebb-and-flow soilless phenotyping platform 15 days after sowing.

|  | RN2 | RN3 | SM | RM | SRL | RTD | RSM | RT | TRL | BD | RD | RV | RA | RLD1 | RLD2 | RLD3 |
| --- | --- | --- | --- | --- | --- | --- | --- | --- | --- | --- | --- | --- | --- | --- | --- | --- |
| RN1 | 0.51 *** | 0.43 *** | 0.35 *** | 0.25 *** | 0.07 ns | -0.07 ns | 0.01 ns | 0.00 ns | 0.29 *** | 0.08 ns | 0.19 * | 0.29 *** | 0.30 *** | 0.28 *** | 0.29 *** | 0.27 *** |
| RN2 | - | 0.55 *** | 0.46 *** | 0.33 *** | 0.07 ns | -0.16 * | 0.03 ns | 0.08 ns | 0.38 *** | 0.09 ns | 0.15 * | 0.36 *** | 0.40 *** | 0.36 *** | 0.37 *** | 0.37 *** |
| RN3 | - | - | 0.24 *** | 0.12 ns | 0.02 ns | -0.03 ns | -0.13 ns | 0.01 ns | 0.13 ns | 0.04 ns | 0.12 ns | 0.25 *** | 0.20 * | 0.10 ns | 0.13 ns | 0.23 *** |
| SM | - | - | - | 0.83 *** | -0.18 * | -0.19 ns | 0.28 ns | 0.33 ns | 0.80 *** | 0.27 ns | 0.21 * | 0.76 *** | 0.84 *** | 0.76 *** | 0.72 *** | 0.81 *** |
| RM | - | - | - | - | -0.30 *** | 0.06 ns | 0.68 *** | 0.33 *** | 0.83 *** | 0.26 *** | 0.25 *** | 0.80 *** | 0.87 *** | 0.78 *** | 0.76 *** | 0.85 *** |
| SRL | - | - | - | - | - | -0.22 *** | -0.25 *** | 0.15 ns | 0.09 ns | 0.29 *** | 0.00 ns | -0.18 * | -0.08 ns | 0.15 * | 0.04 ns | -0.20 *** |
| RTD | - | - | - | - | - | - | 0.06 ns | -0.05 ns | -0.32 *** | -0.05 ns | -0.10 ns | -0.30 * | -0.34 *** | -0.30 *** | -0.31 *** | -0.32 *** |
| RSM | - | - | - | - | - | - | - | 0.20 *** | 0.55 *** | 0.23 *** | 0.29 *** | 0.42 *** | 0.53 *** | 0.53 *** | 0.55 *** | 0.48 *** |
| RT | - | - | - | - | - | - | - | - | 0.47 *** | 0.77 *** | 0.05 ns | 0.21 *** | 0.34 *** | 0.50 *** | 0.38 *** | 0.24 *** |
| TRL | - | - | - | - | - | - | - | - | - | 0.46 *** | 0.32 *** | 0.75 *** | 0.93 *** | 0.99 *** | 0.90 *** | 0.82 *** |
| BD | - | - | - | - | - | - | - | - | - | - | 0.53 *** | 0.16 *** | 0.32 *** | 0.48 *** | 0.51 *** | 0.17 * |
| RD | - | - | - | - | - | - | - | - | - | - | - | 0.25 *** | 0.33 *** | 0.28 *** | 0.56 *** | 0.22 *** |
| RV | - | - | - | - | - | - | - | - | - | - | - | - | 0.93 *** | 0.66 *** | 0.71 *** | 0.96 *** |
| RA | - | - | - | - | - | - | - | - | - | - | - | - | - | 0.87 *** | 0.87 *** | 0.96 *** |
| RLD1 | - | - | - | - | - | - | - | - | - | - | - | - | - | - | 0.86 *** | 0.75 *** |
| RLD2 | - | - | - | - | - | - | - | - | - | - | - | - | - | - | - | 0.72 *** |

Correlation is significant if $p \leq 0.05$ (\*), $p \leq 0.0001$ (\*\*\*) ; ns, no significance (see Table 3 for trait descriptions and units).

### 3.3 Root trait variability

Sixteen root traits and one shoot trait with were included in the PCA. Two principal components (PCs) were identified with eigenvalues >1, capturing 61.7% of the total variation in root traits across the 38 sweetpotato genotypes (Table 6). The first component (PC1) represented 46.2% of the variability and influenced by most root traits (total root length, root mass, root area, root lengths in diameter class 1, 2, and 3, root volume, root shoot mass ratio (Table 6). PC2 represented 15.5% of the total variation and was influenced by the number of roots in node 1, 2, and 3 (Table 6).

**Table 6.** Principal component analysis of 16 root trait and one shoot trait (SM) and the proportion of variation <u>in each principal component.</u>.

| **Traits** | **PC1** | **PC2** |
| --- | --- | --- |
| RN1 | 0.073 | 0.463 |
| RN2 | 0.130 | 0.460 |
| RN3 | 0.061 | 0.496 |
| SM | 0.308 | 0.074 |
| RM | 0.337 | -0.059 |
| SRL | -0.051 | 0.017 |
| RTD | -0.127 | -0.019 |
| RSM | 0.230 | -0.258 |
| RT | 0.088 | -0.333 |
| TRL | 0.340 | -0.076 |
| BD | 0.088 | -0.277 |
| RD | 0.097 | 0.202 |
| RV | 0.327 | 0.061 |
| RA | 0.352 | 0.007 |
| RLD1 | 0.319 | -0.105 |
| RLD2 | 0.331 | -0.022 |
| RLD3 | 0.334 | 0.031 |
| Eigenvalue | 7.85 | 2.63 |
| Variability (%) | 46.2 | 15.5 |
| Cumulative (%) | 46.2 | 61.7 |

The genotype distribution based on the PCA regression scores of the 17 traits (16 root traits and one shoot trait) is shown in a biplot (Figure 5). The biplot shows a well-defined separation of the three root size categories (small, medium, and large root sizes) and a positive contribution of all selected traits except for specific root length and root tissue density (Figure 5). Of the 17 traits, shoot mass, root volume, root area, root mass, total root length and root length by diameter class contributed the most to root size. The relative distance among the 38 sweetpotato genotypes are displayed for each combination of root traits. No evident genotype outliers were observed.

**Figure 5.**
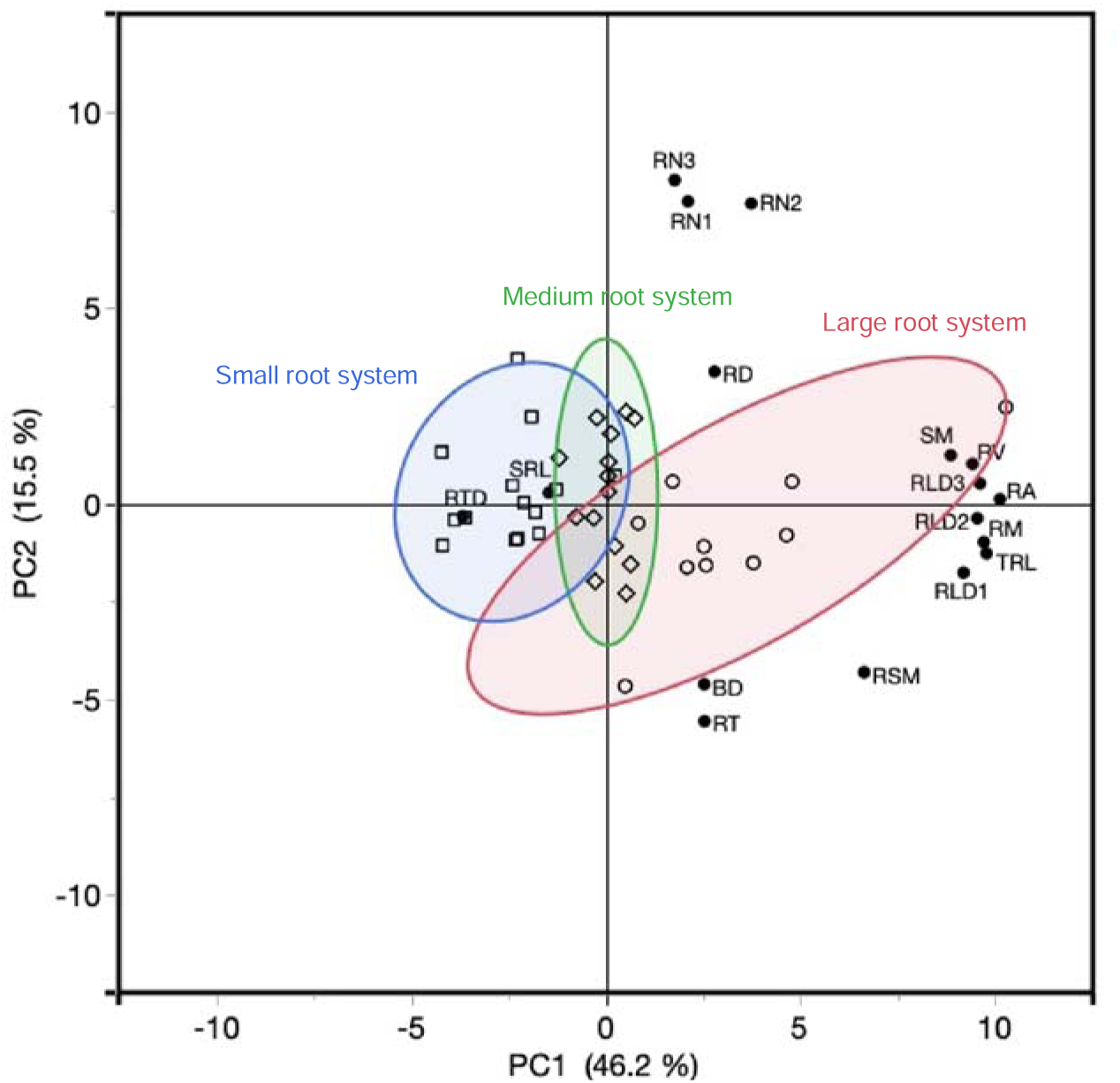
Principal component analysis of 17 traits across 38 sweetpotato genotypes grown in a semi-hydroponic ebb-and-flow soilless phenotyping platform 15 days after sowing. The position of each trait is shown for PC1 vs. PC2 representing 61.7% of the total variability. The genotypes contained in the small root size category are identified with open square symbol and encircled with a blue normal ellipse (*p* < 0.05), the genotypes contained in the medium root size category are identified with open diamond symbol and encircled with a green normal ellipse (*p* < 0.05), and genotypes contained in the large root size category are identified with open circle symbol and encircled with a red normal ellipse (*p* < 0.05).

### 3.4 Genotype clustering based on root trait variation

Hierarchical clustering (constellation plot) of principal component produced four clusters (Figure 6), based on Ward’s criterion that centered cluster creation on minimizing the squared Euclidean distance between points, and indicated quantitative variables associated with each cluster (Figure 6). The 38 sweetpotato genotypes were divided into four Clusters revealing variation in the degree of homogeneity among genotypes examined. Cluster 1 contained 11 genotypes that are characterized by higher values of RTD and lower values of RN1, RN2, SM, RM, RSM, RT, TRL, BD RD, RV, RA, RLD1, RLD2, and RLD3 compared to other genotypes in other clusters (Figure 6). Of these 11 genotypes, all had small root systems. The top four genotypes associated for Cluster 1 are Averre, Mojave, TIS_70683, and Jishu_5. Cluster 2 contained 13 genotypes that are characterized by higher root counts for RN1, and greater SRL and RTD. The following genotypes were representative of Cluster 2: Kakamega, Naspot_8, and NCP09-089. Of these 13 genotypes, one genotype has a large root system, ten had medium root system, and two had small root systems. Cluster 3 (n = 13) contained genotypes with higher RT, BD, RV, RLD1, RLD2, RLD3, SM, RM, RSM compared to Clusters 1 and 2 (Figure 6). The top genotypes with features representative of Cluster 3 were: CN_1483-43, Tinto, NC10-275, and Covington. Cluster 3 revealed eight genotypes with large root systems, four with medium root size system, and one with small root size system. Lastly, Cluster 4 contained one genotype (NCDM04-197; large root system) and was categorized having increased RN1, RN2, RN3, SM, RM, TRL, RD, RV, RA, RLD1, RLD2, and RLD3 compared to all other genotypes and Clusters analyzed (Figure 7). These results determined that genotypes from the same root size were not continually clustered into the same or adjacent group(s).

**Figure 6.**
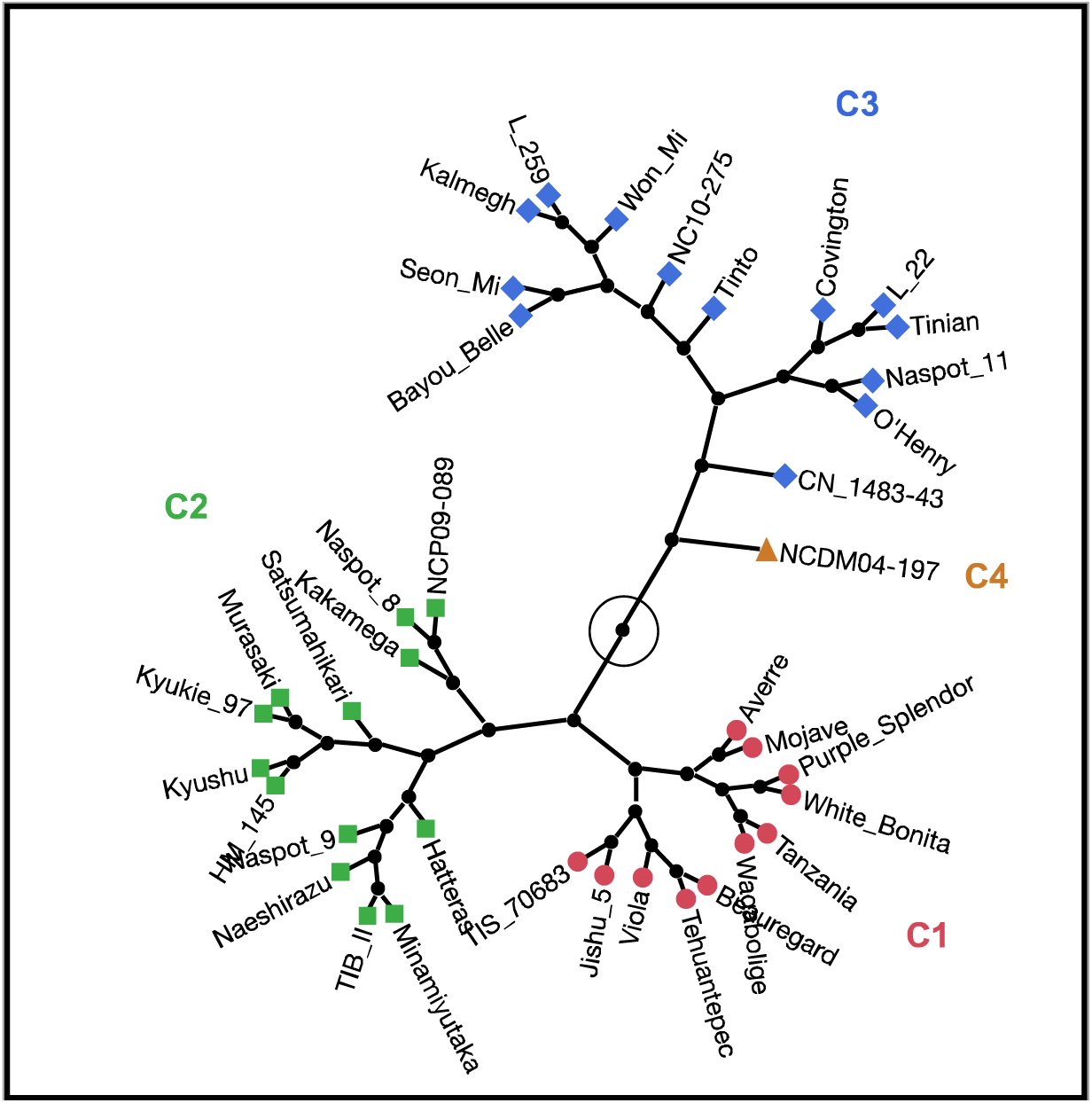
Constellation plot of hierarchical clustering of principal components based on Ward’s criterion method with Euclidean distance as the interval measurement on 16 root traits and one shoot trait. The 38 sweetpotato genotypes were assigned to four clusters (G1 in red, G2 in green, C3 in blue, and C4 in orange).

**Figure 7.**
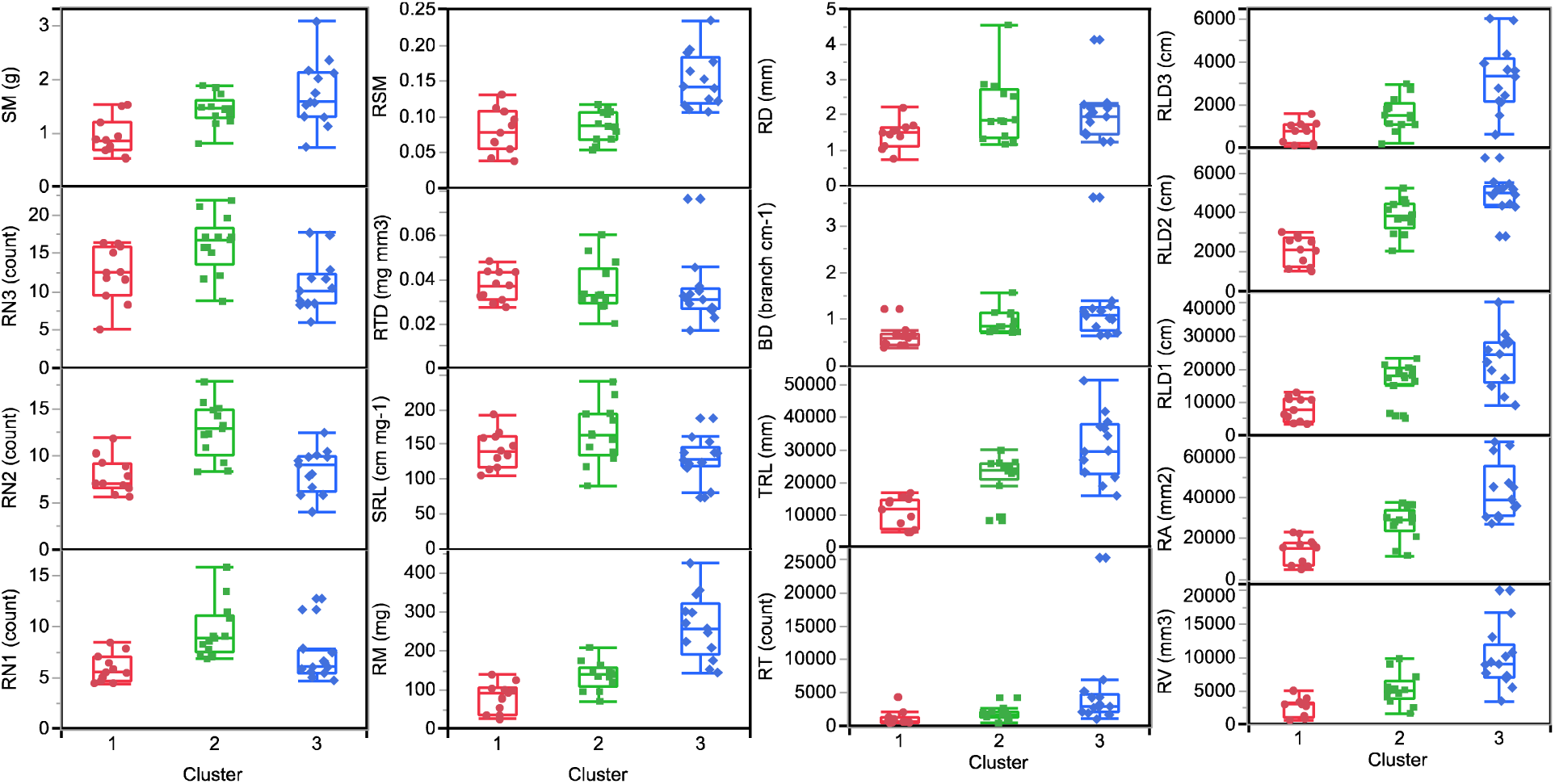
Phenotypic clusters. Box plots show median, 25 and 75 percentiles, and whiskers indicate 1.5 times the interquartile range of the variables assessed within each cluster. The x-axis shows the cluster numbers as presented in Figure 6. Each point represents genotype best linear unbiased predictor. Cluster 4 was omitted as it represents only one genotype.

### 3.5 Correlations in greenhouse root phenotypes related to field performance

Root phenotypes for eight sweetpotato genotypes (e.g., White Bonita, Beauregard, Averre, O’ Henry, Hatteras, Covington, Purple Splendor, and Bayou Belle) were selected and compared to their yield components and performance in a field experiment. Field performance (i.e., TY, MY, and SR#) were associated with SM, RM, TRL, RV and RA (i.e., greenhouse early root traits) (Table 7). Most of the greenhouse root phenotypes did not show a significant correlation with those of harvested plants (∼120 days after planting) in the field. Significant and positive correlations were found between SM, RM, TRL, RV, and RA and field performance (Figure 7). The number of mature storage roots (SR#) was positively correlated with early RM and RV, however there was no significant correlation with SM, TRL, and RA suggesting that TRL, RA, nor SM contribute to this root trait, however these results are only indicative of early root trait utility and suggests additional research to draw a definite conclusion.

**Table 7.** Pearson’s correlation coefficients greenhouse root traits (top row) and field derived phenotypes (leftmost column) in 2022 in Rock Springs, PA.

|  | SM | RM | TRL | RV | RA |
| --- | --- | --- | --- | --- | --- |
| TY | 0.61 *** | 0.59 *** | 0.54 *** | 0.58 *** | 0.58 *** |
| MY | 0.61 *** | 0.58 *** | 0.46 * | 0.57 *** | 0.55 *** |
| SR# | 0.31 ns | 0.41 * | 0.31 ns | 0.40 * | 0.38 ns |

Correlation is significant if $p \leq 0.05$ (\*), $p \leq 0.001$ (\*\*\*); ns, no significance (see Table 3 for trait descriptions and units).

**Figure 7.**
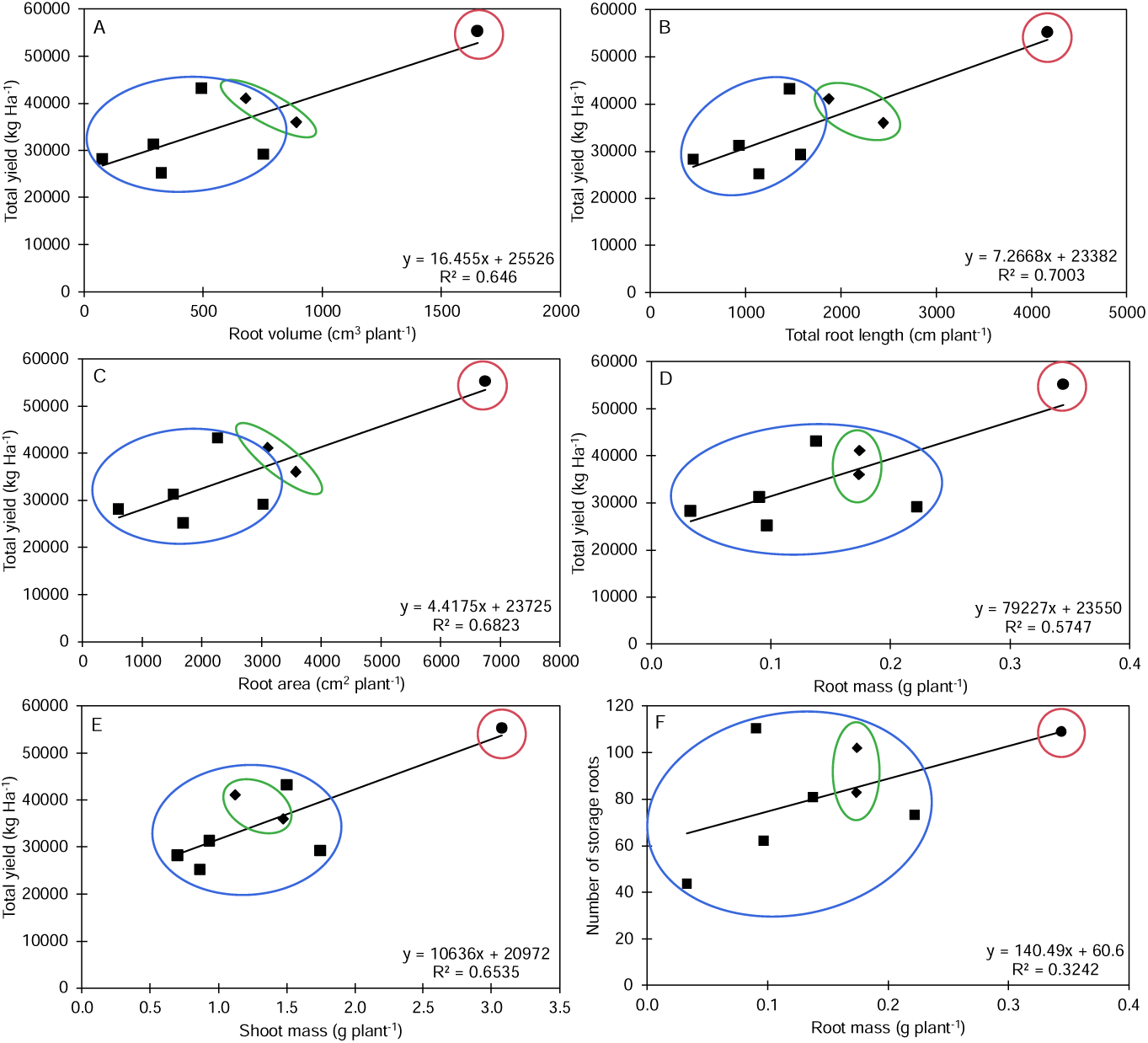
Scatterplots displaying relationships between pairs of phenotypes from greenhouse and field phenotyping. Greenhouse root volume and field total yield (A); greenhouse total root length and field total yield (B); greenhouse root area and field total yield (C); greenhouse root mass and field total yield (D); greenhouse shoot mass and field total yield (E); and greenhouse root mass and field number of storage roots (F). The genotypes contained in the small root size category are identified with square symbol and encircled with a blue normal ellipse, the genotypes contained in the medium root size category are identified with diamond symbol and encircled with a green normal ellipse, and genotypes contained in the large root size category are identified with circle symbol and encircled with a red normal ellipse.

## 4. Discussion

Detecting and optimizing root traits and root system architecture (RSA) by root phenotyping techniques and crop breeding is a crucial factor in enhancing crop production worldwide. However, identifying, assessing, and selecting root traits of diverse sweetpotato germplasm is difficult and it requires innovative techniques that allow for high-throughput and precise quantification of root traits and RSA of sweetpotato throughout their developmental stages. The development of efficient quantitative root phenotyping platforms can improve and aid in the effectiveness and accurateness of collecting genetic data, and measuring phenotypic plasticity due to differing environments.

In this study, we used a semi-hydroponic ebb-and-flow soilless phenotyping platform (Duque, 2022) in combination with a field experiment to characterize a series of root traits in 38 diverse sweetpotato genotypes. Prior studies using a similar system have found that early root traits were linked to variations in yield and nutrient capture [e.g., *Brassica napus* L. (Thomas et al., 2016); *Vigna unguiculata* L. Walp. (Mohammed et al., 2022); *Phaseolus vulgaris* (Strock et al., 2019)], among others. That said, when conducting phenotyping experiments, it is important to consider plant genetic variation. It is well known that plant root systems are very pliable and susceptible to significant G × E effects (Avramova et al., 2016), and the capacity of roots to adjust to different environmental conditions can vary among genotypes and growth environments. In our study, significant correlations were identified between the root traits of 15-day-old sweetpotato slips? cultivated in semi-hydroponic ebb-and-flow soilless phenotyping platform, and field yield. Even though sweetpotato plants grown in an artificial root system phenotyping platform may not exhibit the same development patterns as that of plants cultivated in the field, this approach enabled us to effectively distinguish and analyze the variations in RSA among different sweetpotato genotypes. These findings potentially indicate that early growth RSA of sweetpotato can serve as a reliable predictor of field performance for these agronomic traits.

We measured a total of 20 traits, including 16 early root traits, one shoot trait and three field traits, and identified a large variation among the traits examined in the 38 sweetpotato genotypes. Of these, 14 traits had CVs ≥ 0.5 (Table 4). These individual root trait variations are of particular interest as they could indicate differences in unknown gene functions among the genotypes tested that control root phene state and function. For example, TRL is an important indirect measure of RM (Araki et al., 2002; Valliyodan et al., 2017). In our study, genotypes with the largest root size had four times more TRL and three times more RM compared to genotypes with small root sizes (Figure 2), however no genotypes produced visible root thickening indicative of storage root formation. Also, there was a positive trend between TRL, RM and SM in both greenhouse and field studies (Figures 3, 4, and 7, Table 5 and 7). Root length in diameter class (e.g., RLD1, RLD2, and RLD3) showed significant differences between genotypes contained within the three root sizes (Figure 3C and 4B). These results indicated that RLD1 (fine roots) constitute the bulk of TRL, RM, RV, and RA in sweetpotato. Following this trend, many studies have shown that rapid growth and higher density of finer roots in the topsoil are important for nutrient capture in many crop species (Dunbabin et al., 2003; McCormack et al., 2015; Lynch et al., 2021; Lynch, 2022). Hence, we hypothesize that an early and rapid development of fine roots in sweetpotato can increase its nutrient capture in shallow soils securing carbon allocations for later storage root formation. Indication of strong correlations between SM and RM and several root traits both in the greenhouse as well as in the field, demonstrate that these traits are linked to diverse plant growth approaches (Lynch, 2019; Strock and Lynch, 2020; Villordon et al., 2020; Villordon and Gregorie, 2021).

Many studies indicate that crops exhibiting robust stress tolerance also possess healthy root systems, as well as high rates of respiration in their fine roots. These phenomena likely stem from metabolic activities associated with the absorption and assimilation of nutrients of finer roots (Roumet et al., 2006; Lynch, 2019).

The study’s inclusion of a small yet diverse panel of sweetpotato genotypes suggests that there is a significant level of genotypic variation among them for various root traits, with TRL, RM, RLD, RT, RV, and RA being the most notable ones (as seen in Table 4, Figures 2 and 3). The broad range in these traits observed in the sweetpotato genotypes studied demonstrates a potentially significant level of phenotypic variation for certain soil resource foraging traits, specifically those that are thought to be essential for acquiring nutrients and/or deep-water procurement (Zobel et al., 2007; Lynch, 2019; Wen et al., 2019; Sun et al., 2021). Though sweetpotato is a storage root crop, selection and breeding programs have mainly focused on storage root shape, size, number, and yield, with the impacts on root traits often overlooked (Grüneberg et al., 2015; Mourtala et al., 2023). Beauregard, White Bonita, Purple Splendor, Averre, and O’ Henry ranked in the small size root system category for TRL and RM (Figure 2). Covington, Hatteras, and Murasaki ranked in the medium size root system category, while Bayou Belle ranked in the large root system category (Figure 2). Interestingly, Beauregard (small), Covington (medium), and Bayou Belle (large) are commercially important sweetpotato genotypes with relatively high overall yield and area planted in the United States. These and other recently released sweetpotato genotypes have been bred for high quality, high yielding, and biotic tolerant storage roots under high input field management practices without considering other root and shoot traits. Thus, increased storage root yield through conventional breeding suggests a potential growth and developmental mechanism independent of the size of the root mass early on.

Although sweetpotato is typically recognized as having a shallow root system, having a larger total root length, more lateral roots, and runners (i.e., horizontal stolons with the capacity to “peg down” into the soil and develop roots) can provide the plant with the ability to penetrate deeper into the soil profile and reach water resources that are beyond the reach of shallower-rooted genotypes. The dominance of second-order adventitious roots is evident in the broader range and significantly greater median TRL of sweetpotato root architecture. However, by measuring TRL and other root traits at various depths, or the effect of root formation on trailing vines, which can be substantial, it may be possible to differentiate between various allocation strategies. As mentioned before, our research revealed significant and positive correlations between greenhouse and field root traits (Table 7, Figure 7). These positive correlations between greenhouse-based measurements and field-based measurements of mature root phenotypes demonstrate the effectiveness of using the less labor-intensive and higher throughput greenhouse phenotyping protocol as a valuable tool for breeding programs. Furthermore, phenotyping at an early root developmental stage has proven useful in rapidly characterizing genetic variation for other root phenotypes, such as root hair length and density (Tuberosa et al., 2002; Zhu et al., 2005; Strock et al., 2019). In sweetpotato, Villordon et al. (2020) and Villordon and Gregorie, (2021) showed early species-specific traits regarding root architectural responses to reduced phosphorus and boron availability at the onset of storage root initiation among six sweetpotato cultivars confirming the possibility of early sweetpotato storage root phenotyping.

Although individual root phenotypes may impact performance, enhanced yield under various and changing stress conditions is better understood as the result of interactions among multiple phenotypes that constitute an integrated phenotype (Asfaw et al., 2012; York et al., 2013; Strock et al., 2019). The concept of an integrated phenotype can be evaluated using cluster analysis (constellation plot), offering a valuable new perspective on breeding for edaphic stress tolerance. The uneven distribution of the diversity of sweetpotato, root sizes and genotypes across clusters implies that distinct phene aggregates define gene pools and races (Figures 6 and 7). These separate phene aggregates for each gene pool and race may be associated with adaptive strategies that are specific to the environments from which they originated.

Cluster 1 presented genotypes originating from North America, Far East, Caribbean, Pacific Islands, Africa, and South America. These genotypes were characterized of having reduced and/or smaller root traits compared to all other clusters (Figure 6). Cluster 3 had genotypes from North America, Far East, Pacific Islands, Africa, and Central America with increased and/or larger root traits. Even though root phenotypes could have adverse impacts on performance in the agricultural settings examined in this study, these phenotypes could be better suited to specific edaphic conditions found within the geographic ranges from which these genotypes originated. In contrast to Clusters 1 and 3, Cluster 2 had an intermediate combination of phene states.

Unfortunately, field yield traits were only measured using North American genotypes (more abundant access to genotypes) (Figure 7, Table 7), but our studies showed a positive correlation with several early root traits and yield. Interestingly, genotype Bayou Belle presented a large early root system and the highest field yield. That said, under our research scenario, we did not assess edaphic nutrient stress nor water scarcity for the genotypes herein. Nevertheless, one potential limitation of this system is the premature estimation of the possible field performance of these genotypes. Consequently, under constantly changing growing environments, roots may exhibit specific responses, emphasizing the importance of their inherent phenotypic plasticity for efficient exploitation of edaphic resources (Lynch et al., 2021). Nonetheless, it is still feasible to deduce early genotypic variations among sweetpotato germplasm and their phenotypic plasticity under induced stress treatments.

## 5. Conclusions

This study found significant genetic variation in early adventitious root growth and root system architecture in a small but diverse sweetpotato panel that may relate to differences to local adaptations for both water uptake and nutrient foraging. Selection for TRL and fine roots (e.g., RLD1) at an early stage may be beneficial for sweetpotato grown under edaphic and/or water scarcity. Also, we demonstrate that early RSA can be effectively phenotyped using semi-hydroponic ebb-and-flow soilless phenotyping platform, and that sweetpotato RSA is not necessarily associated with final number of storage roots. Phenotyping sweetpotato for RSA at an early growth stage offers an opportunity to identify genotypes with desired root architecture for improving nutrient foraging and water capture in marginalized soils. Also, this study demonstrated a positive and significant correlation of early root growth with final field yield in a subset of genotypes tested. These specific results suggest that root traits, particularly TRL and RM could improve yield potential and should be incorporated into sweetpotato ideotypes. To help increase sweetpotato performance in challenging environments, breeding efforts may want to consider incorporating early root phenotyping and the idea of integrated root phenotypes.

